# DUET: a graph-based workflow for TCR-epitope prioritization and tumor-reactive T-cell identification

**DOI:** 10.64898/2026.05.27.728217

**Authors:** Yunsheng Chen, Vanessa Giuliano, Ian Dacillo, Wenjun Lin, Yan Yan, Ping Luo

**Affiliations:** Department of Computer Science & Mathematics, Faculty of Computer Science and Technology, Algoma University, Brampton, Ontario L6V 1A3, Canada; Marshall McLuhan Secondary School, Toronto, Ontario M5N 3B1, Canada; College of Computational, Mathematical and Physical Sciences, School of Computer Science, University of Guelph, Guelph, Ontario N1G 2W1, Canada

**Keywords:** T-cell receptor, TCR-epitope binding, heterogeneous graph transformer, tumor-reactive T cells, single-cell RNA-seq

## Abstract

Accurate prioritization of T-cell receptor (TCR)-epitope interactions and identification of tumor-reactive T cells are important but difficult steps in immunotherapy-oriented bioinformatics workflows. Existing methods typically address these tasks separately and either model TCR-epitope pairs as independent observations or rely primarily on transcriptomic signatures. In this study, we present DUET (Dual Unified Evaluation of TCR-Epitopes and Tumor-reactive T cells), a graph-based computational workflow that unifies both applications within a single heterogeneous graph framework. The protocol represents TCRs, epitopes, and T cells as typed nodes connected by similarity and association edges, and combines pretrained sequence embeddings with edge-aware graph attention, Laplacian positional encoding, and bidirectional cross-domain attention. Applied to the IEDB and VDJdb benchmarks, DUET achieved AUROC/AUPR values of 0.937/0.922 and 0.992/0.990, respectively, outperforming five state-of-the-art algorithms. In addition, on a single-cell RNA-seq dataset, the workflow achieved an AUROC of 0.984 and an AUPR of 0.984, substantially exceeding transcriptomic signature-based baselines for tumor-reactive T-cell identification. Ablation analysis showed that Laplacian positional encoding provided the largest performance gain, particularly in sparse graph settings. These results suggest that heterogeneous graph modeling can serve as a practical protocol for integrating receptor sequence, antigen context, and cellular phenotype in computational immunology.

## Introduction

T-cell receptor (TCR) recognition of peptide antigens is central to adaptive immunity and underlies several immunotherapy strategies, including adoptive tumor-infiltrating lymphocyte transfer, TCR-engineered T-cell therapy, and neoantigen vaccine development [1]. However, in practice, two closely related computational problems remain difficult to solve at scale: prioritizing TCR-epitope pairs for downstream validation and identifying tumor-reactive T cells (TRT) from single-cell datasets without relying exclusively on labor-intensive functional assays. Public resources such as IEDB [2] and VDJdb [3] provide growing catalogs of validated TCR-epitope associations, while single-cell studies now jointly profile TCR sequence and cell state in tumor-infiltrating lymphocytes. What is still missing is a practical and reproducible workflow that can integrate these data types in a way that reflects the relational structure of immune repertoires rather than treating each observation in isolation.

Most existing computational approaches address only one part of this workflow. Earlier TCR specificity methods relied on sequence alignment, physicochemical scoring, or conventional machine learning over hand-crafted CDR3 features. Subsequent deep learning approaches, including ERGO-II [4], NetTCR-2.0 [5], DeepTCR [6], and PanPep [7], improved sequence representation learning but generally model each TCR-epitope pair as an independent prediction problem. This limits their ability to exploit repertoire-level structure, where similar TCRs may share binding preferences and related epitopes may recruit overlapping receptor sets. More recent graph-based methods such as GTE [8] and HeteroTCR [9] begin to address this issue for TCR-epitope prediction, but they remain focused on β-chain-centered binding tasks and do not extend naturally to TRT identification from single-cell data. In parallel, current TRT workflows are dominated by transcriptomic signature scoring [10], [11], [12], [13], with relatively limited use of TCR sequence context. As a result, users lack a single computational protocol that can connect receptor sequence, antigen association, and cellular phenotype in one analysis framework.

DUET is a graph-based computational workflow for two related use cases: TCR-epitope binding prioritization and tumor-reactive T-cell identification. The workflow represents TCRs, epitopes, and cells as nodes in a heterogeneous graph linked by intra-domain similarity edges and inter-domain association edges. Sequence-derived features are combined with edge-aware graph attention, Laplacian positional encoding (LPE) [14], and bidirectional cross-domain attention to propagate information within and across biological domains. This design enables the same core protocol to be applied to curated TCR-antigen databases and to single-cell datasets in which TCR clonotypes can be linked to transcriptional cell states. The workflow is evaluated on the IEDB dataset (8,187 TCR-epitope pairs), the VDJdb dataset (71,686 pairs), and a gastrointestinal cancer single-cell dataset (376 functionally labeled T cells [15]). Ablation analysis is performed to determine which structural components contribute most strongly to predictive performance.

## Materials and Methods

### Overview of the DUET Workflow

DUET is a graph-based computational workflow designed for two related applications, prioritizing TCR-epitope associations and identifying tumor-reactive T cells from single-cell RNA-seq data (Fig. 1). In both settings, the workflow operates on a heterogeneous graph in which biologically distinct entities are represented as separate node types connected by similarity-based or association-based edges. The overall procedure consists of four stages, including data preparation, graph construction, feature encoding and graph-based representation learning, and task-specific prediction.

**Figure 1.**
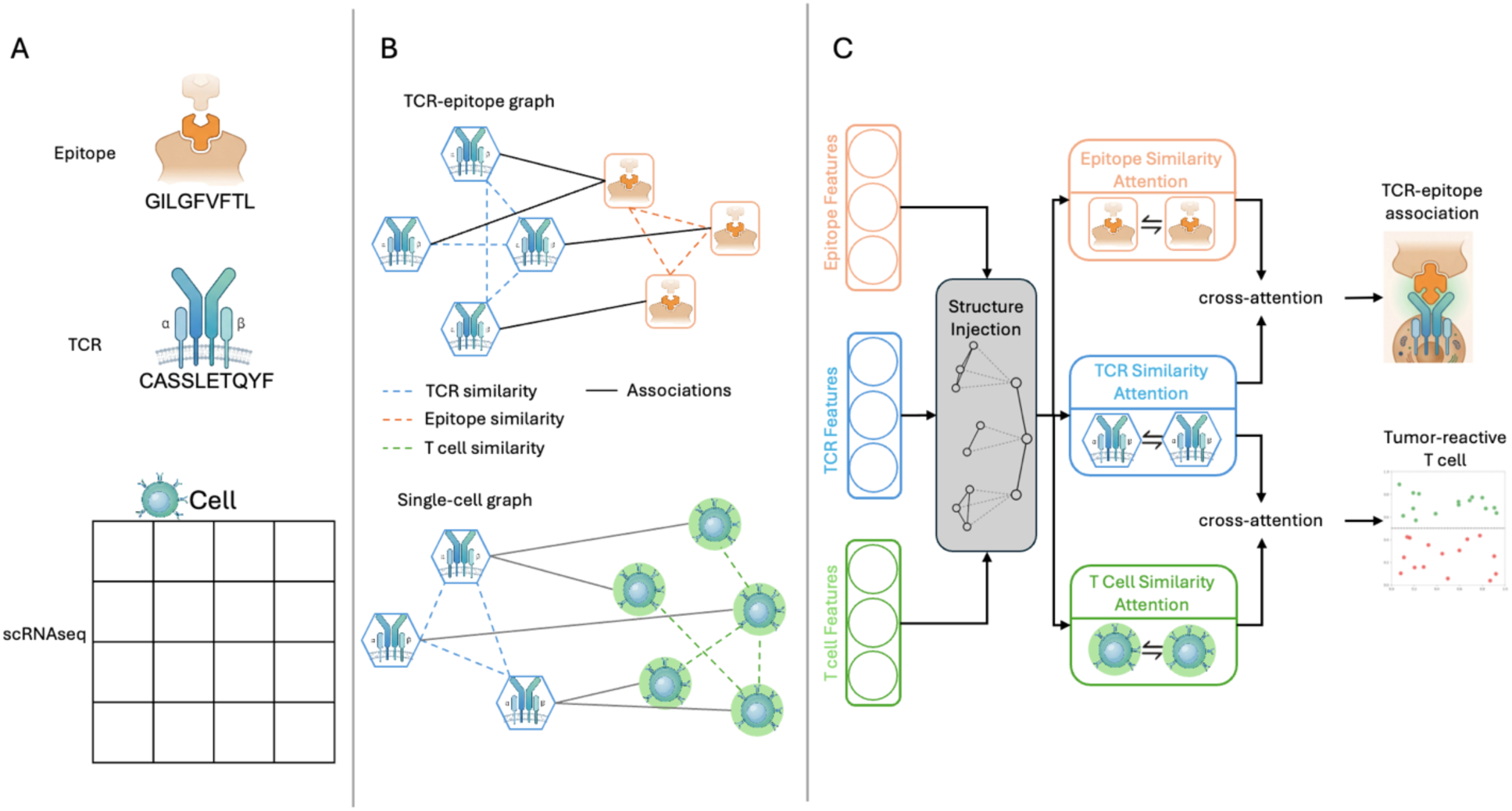
Overview of the DUET workflow. **(a)** Input data types: epitope sequences, TCR CDR3 sequences, and single-cell RNA-seq profiles. **(b)** Heterogeneous graph construction for the TCR-epitope task (top) and the single-cell task (bottom). Nodes represent TCRs, epitopes, or cells; dashed edges denote intra-domain similarity connections; solid edges denote inter-domain associations or clonotype membership. **(c)** Model architecture. Node features and structural encodings are injected via the Structure Injection module, followed by intra-domain graph attention within each node type and bidirectional cross-domain attention between domains. The resulting representations are used for TCR-epitope association prediction (edge-level) and tumor-reactive T-cell identification (node-level).

For TCR-epitope analysis, DUET takes curated TCR sequences and epitope sequences as input and returns a predicted binding score for each candidate TCR-epitope pair. For tumor-reactive T-cell analysis, DUET takes paired TCR and single-cell expression data as input and returns a predicted reactivity score for each cell. The same underlying workflow is used in both applications, with the main differences arising from the input node types and the prediction target.

### Benchmark Datasets

Two public datasets were used to evaluate DUET for TCR-epitope prioritization. The IEDB dataset (version 3) comprised 8,187 curated TCR-epitope pairs after sequence filtering, corresponding to 7,682 unique TCRs and 980 unique epitopes. The VDJdb dataset (version 2025-12-29) comprised 71,686 curated pairs, representing 69,337 unique TCR α- and β-chain CDR3 sequences and 1,352 unique epitopes.

For TRT identification, the gastrointestinal cancer dataset from Zheng et al. (Zheng2022) [15] was used. Following preprocessing and filtering for experimentally validated tumor-reactivity labels and complete paired α- and β-chain CDR3 sequences, the final dataset contained 376 cells, including 62 tumor-reactive and 314 non-reactive cells, representing 84 unique TCR clonotypes.

### Data Preparation

For the IEDB and VDJdb benchmarks, only entries corresponding to *Homo sapiens* and MHC class I were retained. Records were further filtered to include only those with complete curated α- and β-chain CDR3 sequences. Epitope sequences were restricted to canonical peptides composed of the 20 standard amino acids and ranging from 8 to 15 residues in length. TCR CDR3 sequences were further filtered to remove entries containing stop codons, and only sequences 8-20 residues in length were retained for both chains.

Raw count matrices from the Zheng2022 dataset were processed using Seurat v5 [16]. Quality-control filtering retained cells with 200-6,000 detected genes and mitochondrial transcript fractions below 15%. This data was normalized using SCTransform, which allowed highly variable genes to be subsequently identified. Differential expression analysis between tumor-reactive and non-reactive cells was then performed using the Wilcoxon rank-sum test, and significantly differential genes (adjusted p-values below 0.05) were used to derive the cell-level input features for the DUET workflow.

For TCR-epitope prediction, negative examples were defined from unobserved entries of the TCR-epitope association matrix. These negative edges were used only for supervised training and evaluation and were kept separate from held-out positive associations during cross-validation.

### Heterogeneous Graph Construction

DUET represents each dataset as a heterogeneous graph with typed nodes and edges. For the binding-prioritization task, the graph contains TCR nodes and epitope nodes. Inter-domain edges represent known TCR-epitope associations. Intra-domain edges are constructed separately among TCRs and among epitopes using K-nearest-neighbor (KNN) graphs derived from pairwise similarity.

TCR similarity is computed as a combination of k-mer-based cosine similarity and physicochemical similarity. Specifically, sequence-composition similarity was calculated from concatenated bigram (k = 2) and trigram (k = 3) frequency vectors, whereas physicochemical similarity was derived from five amino acid property scales from AAindex [17], including hydrophobicity, normalized net charge, polarity, molecular weight, and isoelectric point. Pairwise cosine similarity was computed for both representations, and the resulting matrices were standardized before integration. For paired TCRs, α- and β-chain similarity matrices were combined as:

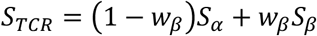

where *w_β_* = 0.7 to reflect the greater contribution of the β chain to antigen specificity. The final TCR similarity matrix was then obtained by averaging the standardized k-mer-based and physicochemical similarity matrices. Epitope similarity is computed analogously, without chain weighting.

For the single-cell task, the graph contains TCR nodes and cell nodes. A cell is connected to the TCR clonotype it expresses through a membership edge. Additional KNN edges are constructed among TCRs based on sequence similarity and among cells based on expression similarity. This graph design allows DUET to capture both direct biological associations and neighborhood structure within each modality.

### Node Feature Initialization

DUET initializes each node type with domain-specific features before graph-based learning. TCR sequences are encoded using BertTCR [18], which is a BERT–based protein language model that generates contextualized embeddings for CDR3 sequences. α- and β-chain CDR3 embeddings are generated separately and concatenated to produce the TCR feature vector. Epitope sequences are encoded using the ESM-2 protein language model [19]. ESM-2 is a transformer-based masked language model trained on millions of protein sequences to capture deep evolutionary, structural, and functional representations of amino acids. Cell nodes are initialized using expression features derived from the differentially expressed genes identified during the single-cell preprocessing workflow.

Before message passing, node features are augmented with structural information. Specifically, DUET incorporates LPE to capture graph-topological context, and similarity edge attributes to weight neighborhood aggregation by pairwise sequence similarity. The augmented node representation is defined as:

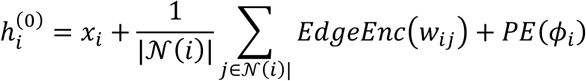

where *x_i_* denotes the initial node feature, *w_ij_* is the edge weight between nodes *i* and its neighbor *j*, *EdgeEnc*(⋅) maps the scalar edge weight to an additive bias term, and *PE*(*φ_i_*) is the projected LPE.

### Graph-Based Representation Learning

The core of DUET consists of two complementary operations: intra-domain refinement and inter-domain information exchange.

#### Intra-domain refinement

Within each node type, DUET applies edge-aware graph transformer layers over the similarity graph. This step updates each node representation by aggregating information from its nearest neighbors while also incorporating edge weights directly into the attention mechanism. In practice, this allows more similar TCRs, epitopes, or cells to exert greater influence during representation learning.

For attention head ℎ, the edge-aware attention logit between nodes *i* and *j* is computed as:

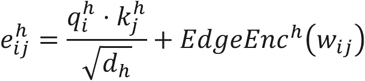

where *q* and *k* are learned query and key projections, *d*. is the per-head hidden dimension. Softmax normalization is then applied across the neighborhood of node *i*.

#### Inter-domain information exchange

After intra-domain refinement, DUET applies bidirectional cross-attention across the relevant biological domains. In the TCR-epitope workflow, this step allows TCR representations to attend to connected epitopes and vice versa. In the single-cell workflow, it allows information exchange between TCR clonotypes and their associated cells. This design enables the model to integrate sequence-level and cell-state information in a shared representation space.

### Task-Specific Prediction

DUET uses the learned graph representations in a task-specific prediction layer. For TCR-epitope prioritization, the final TCR and epitope embeddings for each candidate pair are concatenated and passed to a multilayer perceptron to generate a binding probability. For tumor-reactive T-cell identification, the final cell embeddings are passed to a multilayer perceptron to generate a per-cell tumor-reactivity score. Thus, the same workflow supports both edge-level prediction in the TCR-epitope setting and node-level prediction in the single-cell setting.

### Model Configuration

DUET was implemented in PyTorch [20] and PyTorch Geometric [21], with pretrained sequence embeddings generated using Hugging Face model implementations. Hidden dimension, optimizer, regularization, and training settings were selected separately for the TCR-epitope and single-cell workflows, while preserving the same overall architecture. The final hyperparameter settings used in this study are summarized below, whereas hyperparameter tuning procedures, candidate search ranges, and selection results are provided in the Supplementary Materials.

**Table 1.**
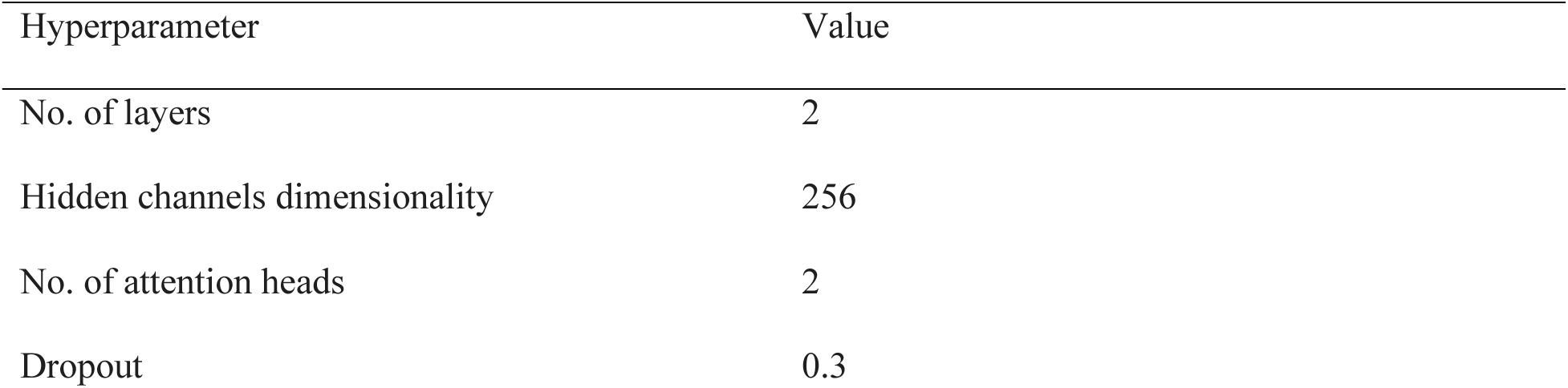
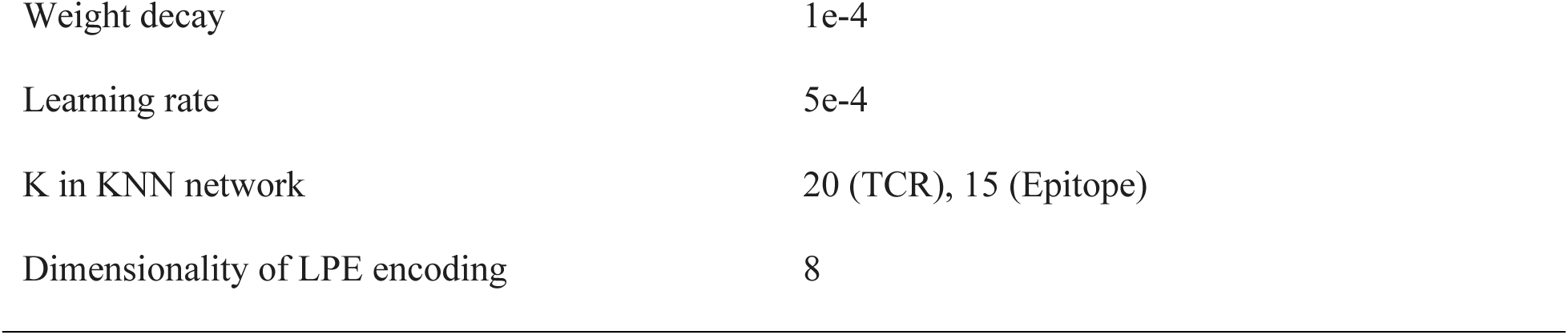
Hyperparameter settings used for DUET.

### Model Evaluation

DUET was evaluated using five repeated runs of 10-fold cross-validation across the three benchmark datasets. Predictive accuracy was assessed using the area under the receiver operating characteristic (ROC) curve (AUROC) and the area under the precision-recall (PR) curve (AUPR). In each fold, known positive associations were partitioned into training (90%) and validation (10%) subsets, and an equal number of negative edges were sampled for each split. To prevent data leakage, inter-domain association edges used during message passing were restricted to positive edges from the training folds only. In contrast, validation-fold edges were excluded from the graph and used solely for held-out evaluation. For DUET, ROC and PR curves were plotted as mean curves across the repeated cross-validation runs, with shaded regions indicating the 95% confidence interval (CI) estimated from run-to-run variability.

For TCR-epitope prediction, DUET was compared with five state-of-the-art deep learning methods: DeepTCR, NetTCR-2.0, HeteroTCR, GTE, and PanPep. For tumor-reactive T-cell prediction, DUET was compared with four gene signature-based approaches (Hanada [11], Lowery [10], Meng [13], and Yossef [12]). Meng et al. reported two sets of signatures (tumor-reactive and non-tumor-reactive), so both variants were evaluated separately as Meng_TR and Meng_NTR.

To assess generalization to unseen TCR sequences, TCR-disjoint cross-validation was also performed, in which TCR clonotypes in the test set were strictly excluded from training. This protocol evaluates whether DUET retains predictive ability when receptor identities are entirely novel, rather than interpolating within observed sequence space.

## Results

Using the evaluation framework, DUET was next compared with existing baseline methods across the IEDB, VDJdb, and Zheng2022 benchmarks. Across all three datasets, DUET consistently achieved the highest performance in both ROC and PR analyses, with narrow 95% CI across five repeated runs, indicating strong predictive accuracy and stable model behavior.

As shown in Fig. 2, DUET achieved an AUROC of 0.937 (95% CI, 0.931-0.942) and an AUPR of 0.922 (95% CI, 0.909-0.936) on the IEDB benchmark. Among the five baselines, NetTCR-2.0 showed the highest overall performance (AUROC=0.870, AUPR=0.892), followed by HeteroTCR (AUROC=0.814, AUPR=0.843), while GTE, PanPep, and DeepTCR showed lower overall performance. The near-zero AUPR observed for DeepTCR likely reflects a mismatch between its generative architecture and the link- prediction setting, consistent with known limitations of repertoire-based generative models on pairwise association tasks.

**Figure 2.**
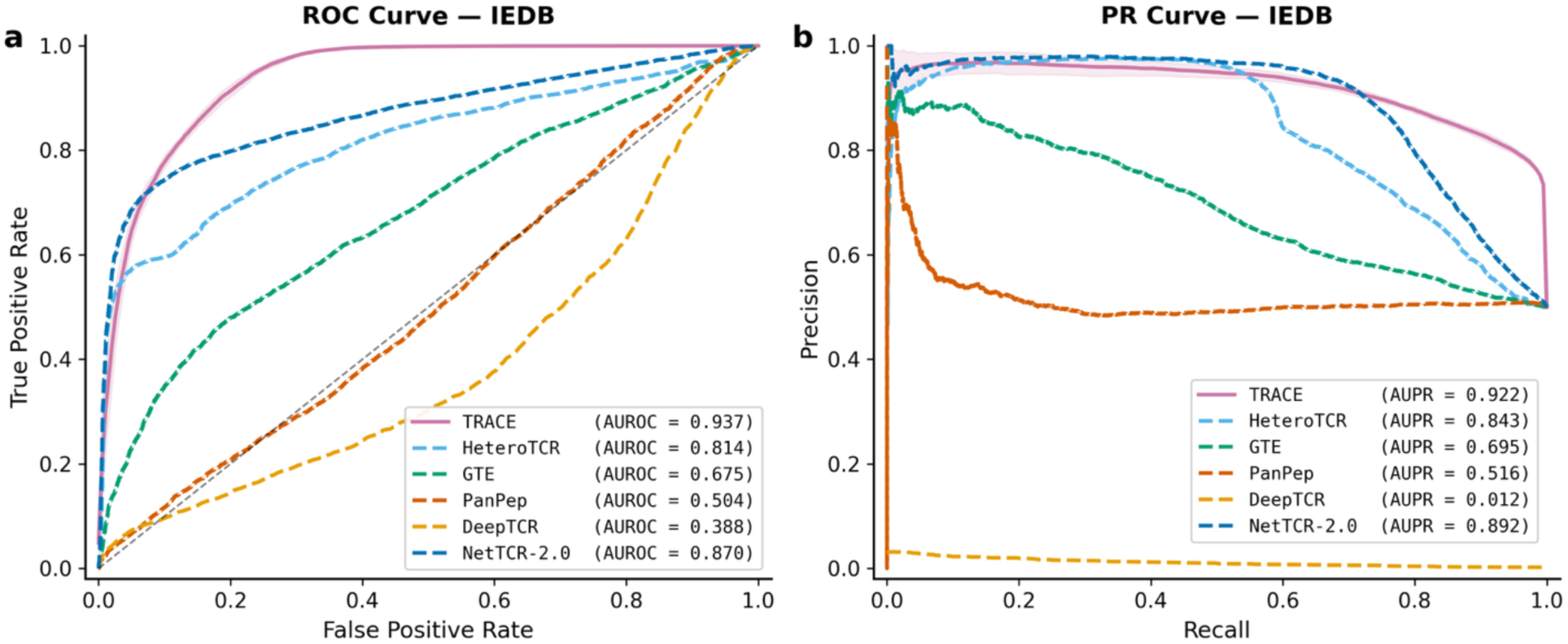
Predictive performance on the IEDB benchmark. **(a)** ROC curves and **(b)** PR curves for DUET and five baseline methods. AUROC and AUPR values are shown in the legend. Curves for DUET represent the mean across five repeated 10-fold cross-validation runs.

A similar trend was observed on the VDJdb benchmark (Fig. 3). DUET achieved an AUROC of 0.992 (95% CI, 0.991-0.993) and an AUPR of 0.990 (95% CI, 0.989-0.991), maintaining the top rank across both metrics. However, on this larger dataset, the top-performing methods converged to similarly high performance. While DUET consistently outperformed competing approaches, the narrower performance margins relative to IEDB likely reflect the denser graph structure of VDJdb, which contains substantially more associations and provides a sufficient training signal for all graph-based methods to converge.

**Figure 3.**
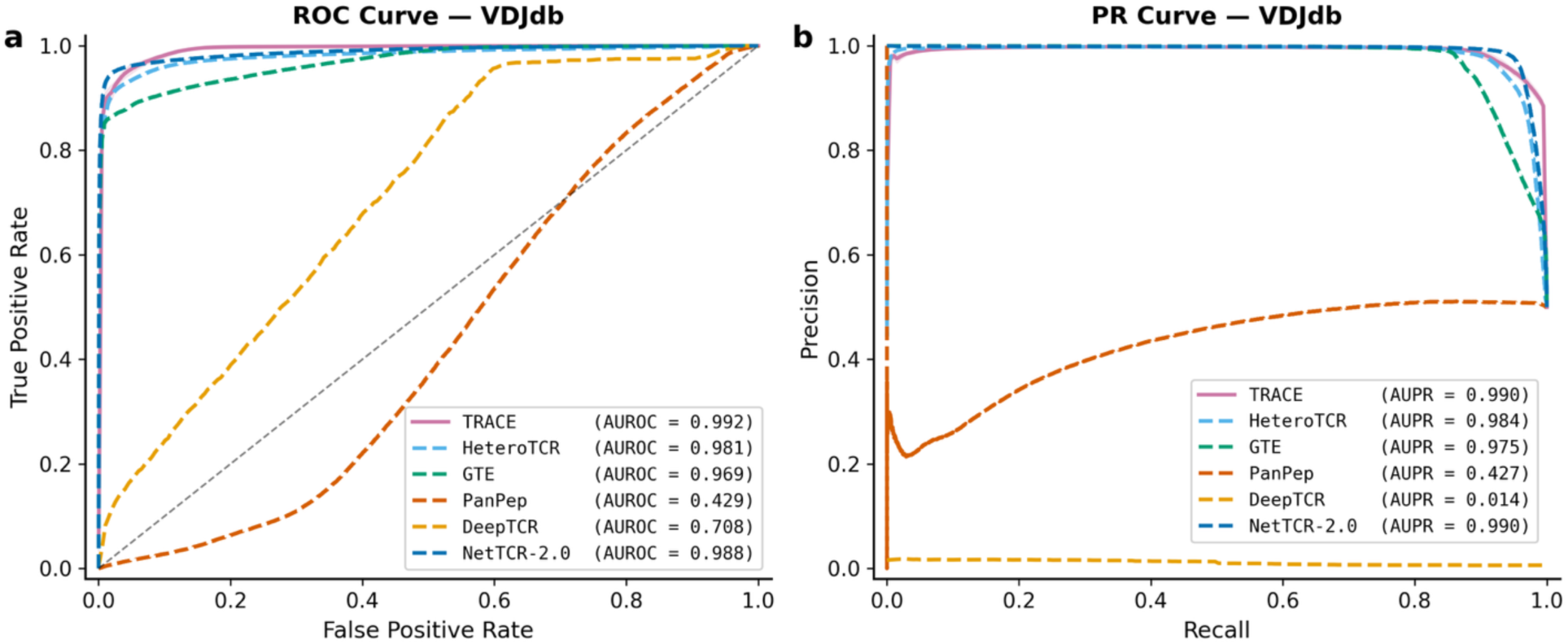
Predictive performance on the VDJdb benchmark. **(a)** ROC curves and **(b)** PR curves for DUET and five baseline methods.

Taken together, these results demonstrate that DUET generalizes robustly across independent TCR–antigen datasets, with the largest gains observed in sparser, more degree-skewed settings where structural encodings are most informative.

On the Zheng2022 scRNAseq benchmark, DUET achieved an AUROC of 0.984 (95% CI, 0.973-0.994) and an AUPR of 0.984 (95% CI, 0.976-0.991), exceeding all evaluated signature-based baselines (Fig. 4). The best competing method, Lowery, obtained an AUROC of 0.735 and an AUPR of 0.310, whereas Hanada, Yossef, Meng_TR30, and Meng_NTR15 all showed substantially weaker discrimination. The AUPR gap is especially pronounced, where DUET’s 0.984 represents a three-fold improvement over the best baseline, indicating that signature-based approaches struggle to maintain precision when recovering the minority tumor-reactive class. These findings suggest that DUET captures tumor-reactivity signals more effectively by jointly modeling TCR sequence, clonotype-level graph structure, and transcriptional cell state, rather than relying on pre-defined gene expression programs.

**Figure 4.**
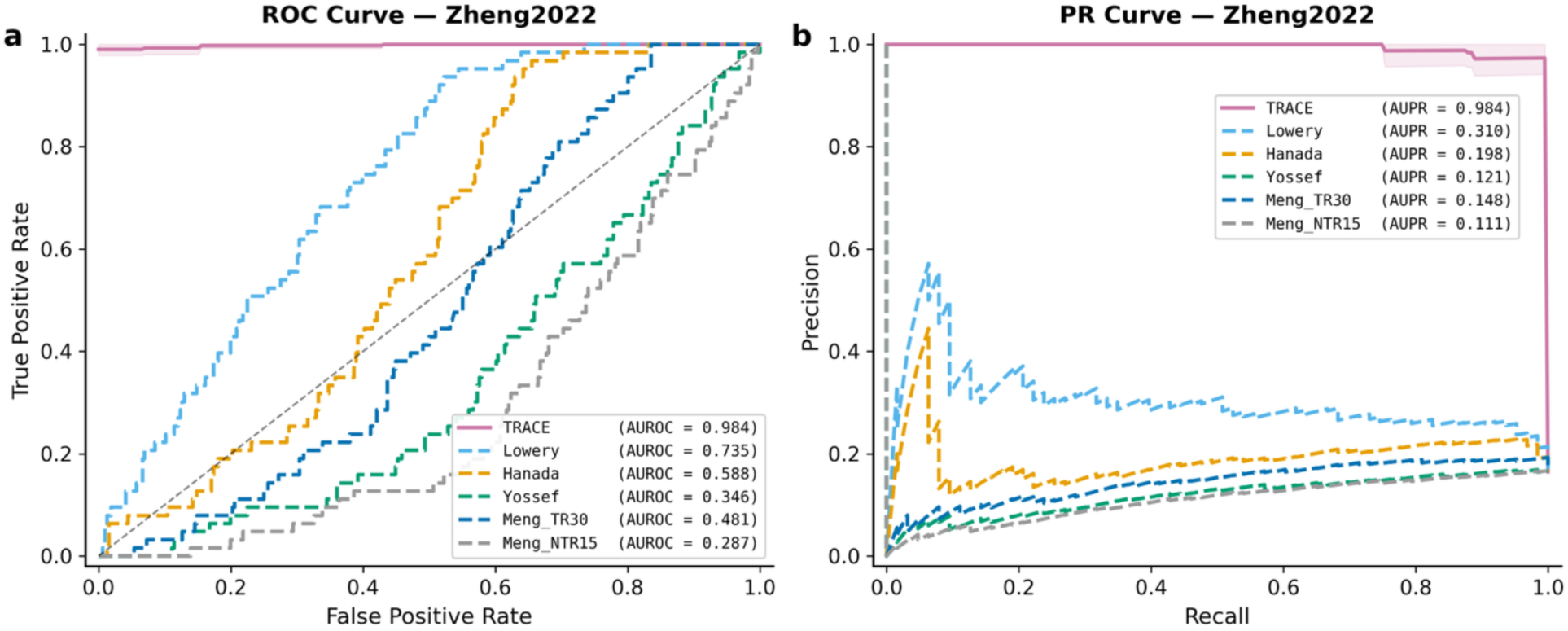
Predictive performance on the Zheng2022 single-cell benchmark. **(a)** ROC curves and **(b)** PR curves for DUET and four transcriptomic signature-based baselines.

These results demonstrate that DUET provides consistently superior predictive performance across both paired TCR-epitope prediction and tumor-reactive T-cell identification. The especially strong gains in AUPR further indicate that the model is well suited to biologically realistic settings in which positive reactive examples are relatively sparse.

### TCR-disjoint Cross-Validation

To evaluate whether DUET generalizes beyond memorization of observed TCR-epitope associations, the evaluation was repeated under TCR-disjoint cross-validation, where test-set TCR sequences were held out entirely from training (Fig. 5). On IEDB, DUET retained strong performance with an AUROC of 0.888 and AUPR of 0.906, representing a modest decline relative to standard cross-validation. On VDJdb, performance was largely preserved (AUROC=0.983, AUPR=0.986), consistent with the dataset’s larger scale and denser graph structure providing sufficient coverage even under sequence-level exclusion. On the Zheng2022 dataset, AUROC remained at 0.893, though AUPR declined to 0.733, likely reflecting the small number of unique clonotypes (n=84) in this dataset, where disjoint splitting substantially reduces training coverage of the TCR neighborhood. These results indicate that DUET generalizes meaningfully to unseen TCR sequences, with the most pronounced sensitivity occurring in the smallest dataset where clonotype-level held-out evaluation is most constrained.

**Figure 5.**
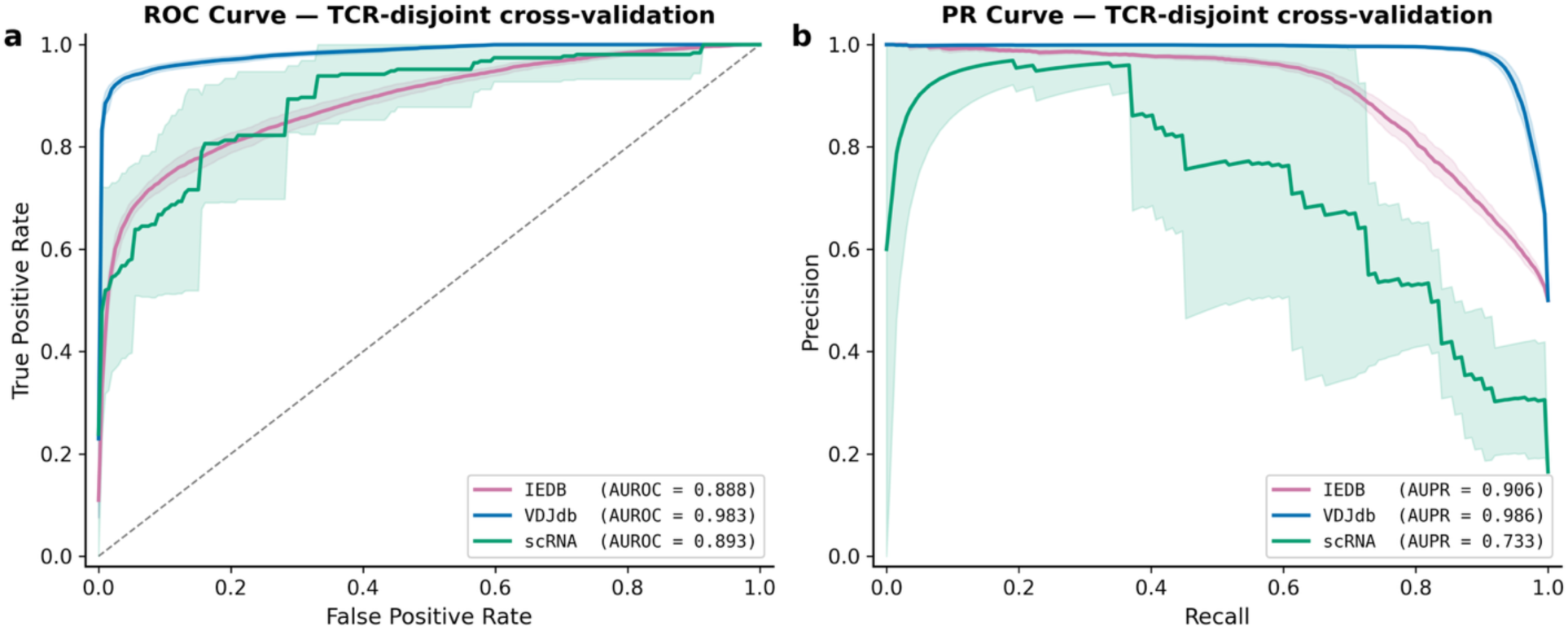
TCR-disjoint cross-validation performance. **(a)** ROC curves and **(b)** PR curves for DUET evaluated on IEDB, VDJdb, and the scRNA dataset under TCR-disjoint cross-validation, in which TCR sequences in the test set were entirely withheld from training. Shaded regions indicate 95% CI across five repeated runs.

### Ablation Study

To isolate the contribution of each structural encoding component, three progressively enriched model variants were evaluated across all three benchmarks. This includes a base model with no structural encoding, the base model augmented with LPE, and the full model that further incorporates similarity edge attributes (Base + LPE + Edge Attributes). Results are shown in Fig. 6.

**Figure 6.**
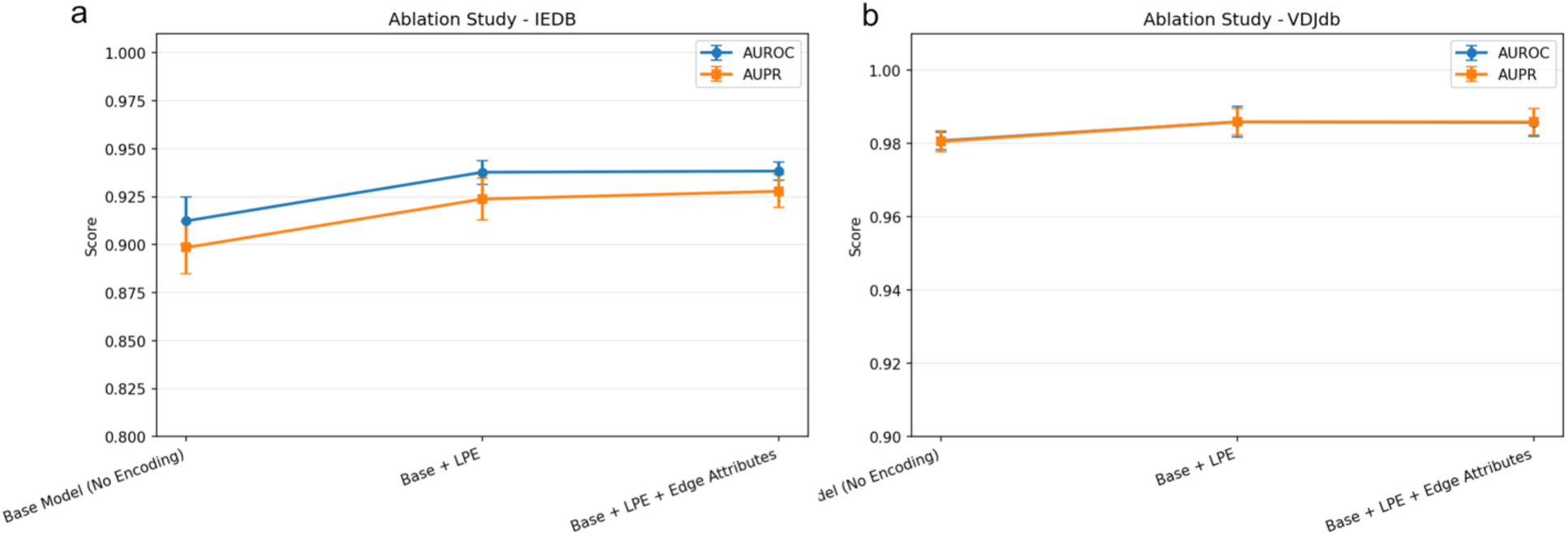
Ablation study results. Mean AUROC and AUPR scores (±95% CI) for three progressively enriched model variants on **(a)** IEDB and **(b)** VDJdb. The three variants are: Base Model (no structural encoding), Base + LPE, and Base + LPE + Edge Attributes.

On the IEDB benchmark (Fig. 6a), removing all structural encodings yielded a base performance of AUROC=0.912 and AUPR=0.899. Adding LPE produced a substantial and consistent improvement, raising AUROC to 0.938 and AUPR to 0.924. This finding indicates that graph-topological context captured by LPE is informative in this setting, likely because the IEDB graph is comparatively sparse and degree-skewed, whereas local spectral structure provides signals that are not recoverable from node features alone. The addition of similarity edge attributes yielded only a marginal increment (AUROC=0.938, AUPR=0.928), suggesting that edge weights contribute modestly once positional structure is encoded.

On the VDJdb benchmark (Fig. 6b), all three variants converged to near-ceiling performance. The base model already achieved an AUROC of 0.979 and AUPR of 0.978, and the addition of LPE and edge attributes produced incremental improvements that remained within the confidence interval range. This near-uniform performance across variants is consistent with the substantially larger and denser graph structure of VDJdb, which contains more TCR-epitope associations and provides sufficient training signal for all model variants to converge regardless of structural encoding.

Notably, degree encoding was also evaluated as an additional structural component. Contrary to expectations, incorporating degree encoding consistently degraded performance across both the IEDB and VDJdb benchmarks (Supplementary Fig. S1, S2). This can be attributed to the highly skewed degree distribution in the TCR-epitope graphs, where hubs may cause degree embeddings to introduce a confounding bias rather than a useful structural signal. Based on these results, degree encoding was excluded from the final DUET architecture.

On the scRNA dataset, all variants achieved saturated performance (AUROC and AUPR of 1, Supplementary Fig. S3), likely due to the small dataset size of 376 cells.

The ablation results suggest that LPE provides the most consequential structural signal among the tested components, particularly in data-scarce or graph-sparse regimes such as IEDB where topological context is otherwise difficult to recover from node features and connectivity alone.

## Discussion

DUET was developed to address two closely related problems in computational immunology: prediction of TCR-epitope binding and identification of tumor-reactive T cells. The results show that a unified heterogeneous graph transformer can perform strongly on both tasks. Across the IEDB and VDJdb benchmarks, DUET consistently outperformed sequence-based and graph-based baselines in both AUROC and AUPR, indicating gains not only in overall discrimination but also in the ability to recover true positives under class imbalance. The model also achieved substantially better performance than transcriptomic signature-based approaches on the Zheng2022 single-cell dataset, suggesting that joint modeling of receptor sequence, graph structure, and cell state captures biologically meaningful information that is missed when each modality is considered in isolation.

These findings support the central design hypothesis that TCR recognition is more effectively modeled within a relational context than as a collection of independent sequence pairs. Earlier deep learning approaches, such as DeepTCR and NetTCR-2.0, improved predictive performance through richer sequence representations but still treated each TCR–epitope pair largely as an isolated instance. Similarly, sequence-based models like NetTCR-2.0 and PanPep encode residue-level features without incorporating broader repertoire context, limiting their ability to exploit shared binding preferences among similar receptors. Although DeepTCR adopts a repertoire-level framework, it is not designed for pairwise link prediction. More recent graph-based methods, including GTE and HeteroTCR [8], [9], partially address these limitations by modeling heterogeneous TCR-epitope interactions. However, they remain primarily focused on binding prediction, rely predominantly on β-chain CDR3 information, and do not extend to single-cell tumor-reactivity tasks. In contrast, DUET integrates intra-domain similarity structure with inter-domain attention, enabling information propagation across related TCRs, epitopes, and, within single-cell settings, phenotypically similar cells. It further incorporates paired α/β chain embeddings where available and generalizes seamlessly to cell-level reactivity prediction without architectural modification. These design choices lead to consistent performance gains, particularly in AUPR, which is a more informative metric than AUROC in settings characterized by sparse positives. The improvement is especially pronounced on the IEDB dataset, underscoring the advantage of leveraging repertoire-level graph context in data-sparse regimes typical of practical immunotherapy applications.

The ablation results clarify which structural components contribute most to model performance. On IEDB, where the graph is sparser and more degree-skewed, LPE provided the largest performance gain. This suggests that graph-topological context captured by spectral positional encoding is particularly informative when training data are limited and connectivity is heterogeneous. The addition of similarity edge attributes yielded a further but modest improvement, indicating that explicit edge-weight information complements but does not substitute for positional encoding. On VDJdb, all variants converged to near-ceiling performance, consistent with the denser graph structure and larger training set providing sufficient signal regardless of structural encoding. We interpret the limited impact of edge attributes cautiously as a dataset-dependent result rather than evidence of general uninformativeness, and note that larger and more structurally heterogeneous datasets will be needed to fully characterize the contribution of edge-level encodings. In the single-cell dataset, all ablated variants similarly remained near ceiling, which is consistent with the small dataset size and limited clonotype diversity, suggesting that larger single-cell cohorts will be needed to fully resolve the contribution of each structural module.

The TCR-disjoint cross-validation results further support generalizability on larger datasets. However, the more pronounced AUPR decline on the small scRNA cohort underscores that disjoint evaluation on datasets with limited clonotype diversity will require larger single-cell cohorts to be fully characterized.

Several limitations should be considered when interpreting these results. First, the scRNA analysis was conducted on a relatively small dataset from a single study [15]. Although the model outperformed published signature-based baselines, the observed performance may overestimate generalizability to other cancer types, treatment settings, or sequencing platforms. At present, Zheng2022 is the only TRT single-cell dataset available for our evaluation, reflecting the limited public availability of comparable datasets under privacy and controlled-access constraints. To address this limitation, access to additional datasets from studies [10], [11], [12], [13] are being sought for future external validation of the model. Second, as in many TCR specificity studies, negative TCR-epitope pairs are inferred from unobserved combinations rather than experimentally verified non-binders. This benchmarking strategy is practical but imperfect as some presumed negatives may be untested positives.

Despite these limitations, the present study highlights a promising direction for bioinformatics methods in adaptive immunity. A model that can jointly learn from receptor sequence, antigen context, and single-cell phenotype has clear relevance for TCR discovery, prioritization of tumor-reactive clonotypes, and the broader design of personalized immunotherapies. Future work should focus on expanding evaluation to larger multi-patient cohorts, incorporating explicit MHC information, and improving interpretability so that the graph signals driving each prediction can be linked back to biological mechanisms. More broadly, the results suggest that heterogeneous graph learning provides a flexible framework for integrating repertoire, antigen, and cellular data, and may serve as a foundation for next-generation predictive models in computational immunology.

## Conclusions

DUET provides a graph-based computational workflow for two related immunological tasks, namely prioritizing TCR-epitope associations and identifying tumor-reactive T cells from single-cell data. By integrating sequence-derived representations with similarity-aware graph structure and cross-domain attention, the protocol captures both molecular and repertoire-level context in a unified framework. Across the IEDB, VDJdb, and Zheng2022 benchmarks, DUET showed strong and consistent predictive performance, with particularly favorable precision-recall behavior relative to competing approaches. Ablation analysis further indicated that LPE is the most consequential structural component, particularly in sparse and degree-skewed graph settings where topological context is otherwise difficult to recover from node features alone. Taken together, these results support heterogeneous graph modeling as a practical strategy for linking TCR sequence, antigen specificity, and cellular phenotype, positioning DUET as a useful protocol for bioinformatics workflows in immunotherapy and T-cell repertoire analysis.

## Key Points

● DUET is a heterogeneous graph-based workflow that unifies TCR-epitope binding prioritization and tumor-reactive T cell identification within a single computational framework.
● On the IEDB and VDJdb benchmarks, DUET outperformed five state-of-the-art methods, achieving AUROC/AUPR of 0.937/0.922 and 0.992/0.990 respectively.
● On a gastrointestinal cancer single cell dataset, DUET achieved an AUPR of 0.984, nearly three times higher than the best transcriptomic signature-based baseline.
● Ablation analysis demonstrated that Laplacian positional encoding contributes most meaningfully in sparse, degree-skewed graph settings, while all variants converged under denser graph conditions.
● TCR-disjoint cross-validation confirmed that DUET generalizes to unseen TCR sequences, supporting its applicability to novel receptor repertoires beyond the training data.

## Acknowledgements

This work was supported by the Algoma University Research Fund; and the Natural Sciences and Engineering Research Council of Canada (NSERC) [grant number: RGPIN-2025-06236].

## Conflict of Interest

The authors declare no conflict of interest.

## Data Availability

The code and data will be made publicly available on GitHub upon acceptance.

## Author Biography

Ping Luo is an Assistant Professor in the Department of Computer Science & Mathematics at Algoma University, Canada. His research focuses on computational biology, machine learning, and bioinformatics applications in next generation sequencing.

## Notes

### Competing Interest Statement

The authors have declared no competing interest.

### Summary of Updates

Title and name of the framework.

