## Supplementary Methods for "DUET: a graph-based workflow for TCR-epitope prioritization and tumor-reactive T-cell identification"

### Supplementary Material

#### Supplementary Methods

Hyperparameter tuning was performed on the IEDB benchmark using a grid search over candidate values for parameter in Table S1. The tuned hyperparameters included the number of graph transformer layers, hidden channel dimensionality, the number of attention heads, and the number of nearest neighbors (K) used in KNN graph construction. Other hyperparameters, such as dropout rate, weight decay, learning rate, and the dimensionality of the Laplacian positional encoding were chosen based on experience.

Candidate search ranges for each hyperparameter are also provided in Table S1. The combination yielding the highest mean AUROC runs on IEDB was selected as the final configuration. This same hyperparameter set was then applied without further modification to the VDJdb and Zheng2022 benchmarks, allowing performance differences across datasets to be attributed to dataset characteristics rather than dataset-specific tuning.

#### Supplementary Table S1

| Hyperparameter | Ranges |
|---|---|
| No. of graph transformer layers | 1, 2, 3 |
| Hidden channel dimensionality | 64, 128, 256 |
| No. of attention heads | 2, 4, 8 |
| No. of nearest neighbors | 10, 15, 20, 25 |

### Supplementary Figures

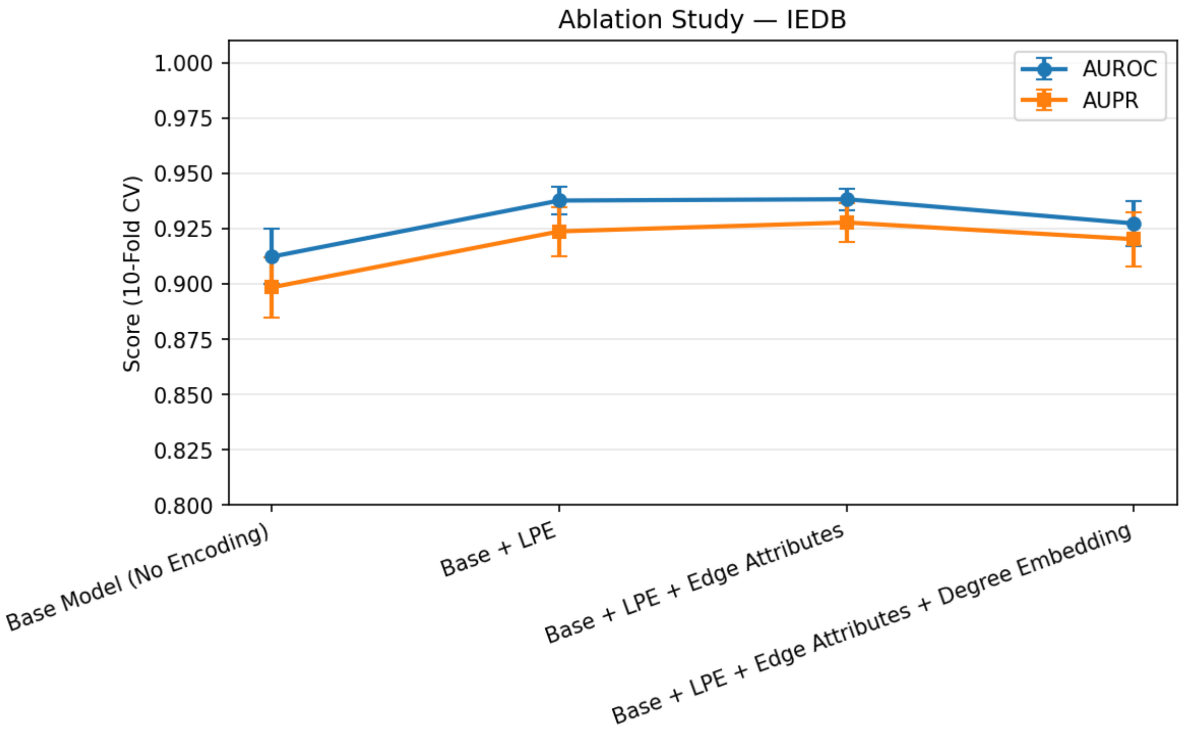


Figure S1. Ablation Study with Degree Encoding (IEDB)

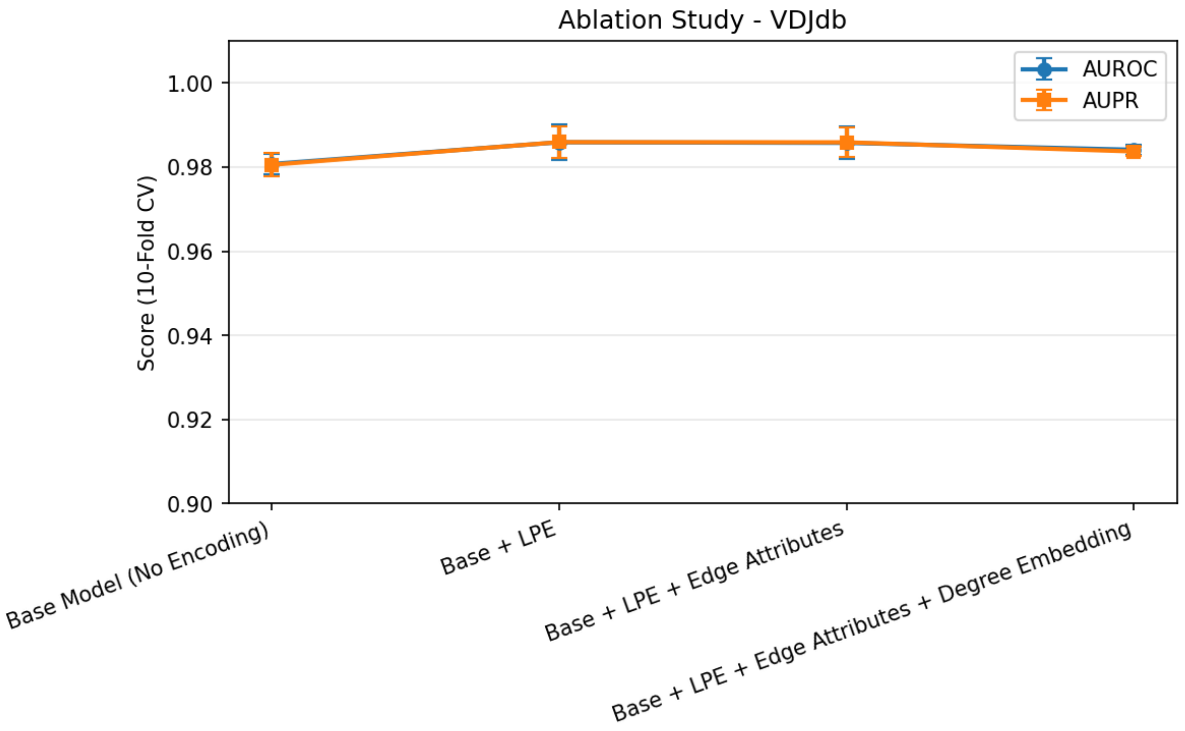


Figure S2. Ablation Study with Degree Encoding (VDJdb)

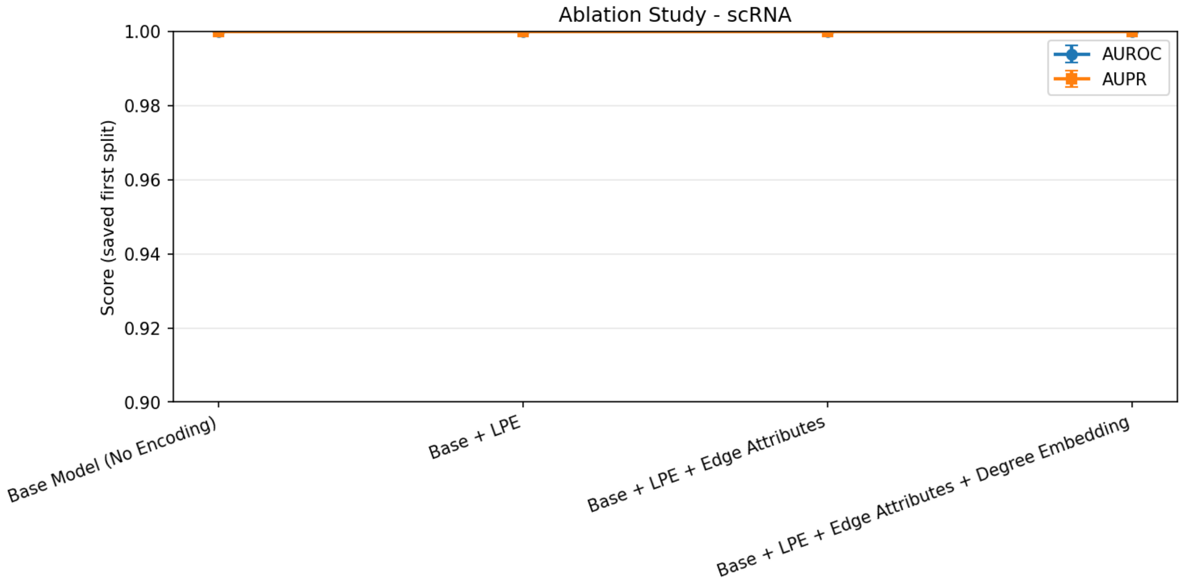


Figure S3. Ablation Study with Degree Encoding (Zheng2022)
